# Complex Modulation of IL-6 Signaling by Apelin and Elabela in HTR-8/SVneo Cells Under Cobalt Chloride-Induced Chemical Hypoxia

**DOI:** 10.64898/2026.08.20.746041

**Authors:** Anna J. Soloshenko, Courtney Brown, Xuming Sun, Ajay N. Roy, Jonathan Ray, Ebrahim Elsangeedy, Mark C. Chappell, Liliya M. Yamaleyeva

## Abstract

Preeclampsia is a pregnancy complication characterized by hypertension, proteinuria, and end-organ dysfunction. Abnormal placentation leading to reduced placental perfusion may contribute to its development. Previous studies demonstrated that the activation of the apelin receptor (APJ) system has hypotensive, renoprotective, and antioxidant effects in preeclamptic rat models. Apelin and elabela (ELA) can stimulate the proliferation of trophoblast cells, suggesting a role in embryonic development. However, the mechanisms underlying the actions of apelin or ELA in trophoblast cells are not well understood, particularly in hypoxic settings. The immortalized HTR-8/SVneo trophoblastic cells were treated with cobalt chloride (CoCl_2_) at 0.2 mM for 24 hours to mimic hypoxic conditions. RT-qPCR, ELISA or Western blotting was used to measure mRNA or protein levels of apelin, elabela, and the components of IL-6 signaling in cell lysates or conditioned media. The exposure to CoCl_2_ increased total apelin and elabela content approximately 2-fold in the conditioned media but did not affect APJ levels. CoCl_2_ upregulated proinflammatory cytokine concentrations: soluble fms-like tyrosine kinase 1 (sFlt-1), soluble gp130 (sgp130), interleukin-6 (IL-6), and sIL-6 receptor (IL-s6R). Both apelin and elabela downregulated IL-6 mRNA but had no effect on sFlt-1 mRNA. Apelin attenuated sgp130, while ELA decreased the membrane form of IL- s6R. Apelin also decreased the pSTAT3/STAT3 ratio. CoCl_2_-induced hypoxia upregulated the pro-inflammatory milieu in HTR-8/SVneo cells. Local activation of this peptidergic system may be a compensatory response of the trophoblast cells to hypoxia as exogenous apelin and elabela treatment ameliorated the hypoxia-induced pro-inflammatory milieu.

**Highlights:**

- ur data confirmed that both membrane and soluble forms of IL-6R, and the components of IL-6 trans-signaling such as soluble and membrane-bound gp130 are present in HTR-8/SVneo trophoblastic cells.
- CoCl_2_ exposure upregulated apelin and elabela but not apelin receptor APJ in HTR-8/SVneo cells.
- IL-6 and soluble gp130 were upregulated by CoCl_2_ exposure which was reversed by apelin or elabela. Elabela also downregulated membrane-bound IL-6R in CoCl_2_-exposed HTR-8/SVneo cells.
- IL-6 signaling through its soluble receptor primarily drives inflammation, whereas signaling via the membrane-bound receptor tends to support cell survival and anti-inflammatory effects, our data suggest that apelin or elabela could be involved in the complex regulation of the hypoxic environment in trophoblast cells relevant for preeclampsia.

**Graphical abstract:** Schematic of proposed interactions between IL-6 signaling and apelin and elabela in HTR-8/SVneo trophoblastic cells under CoCl₂-mimicked hypoxia. We hypothesize that apelin/elabela/APJ axis regulates IL-6 signaling via its actions on various components of the IL-6 classical and trans-signaling pathway.

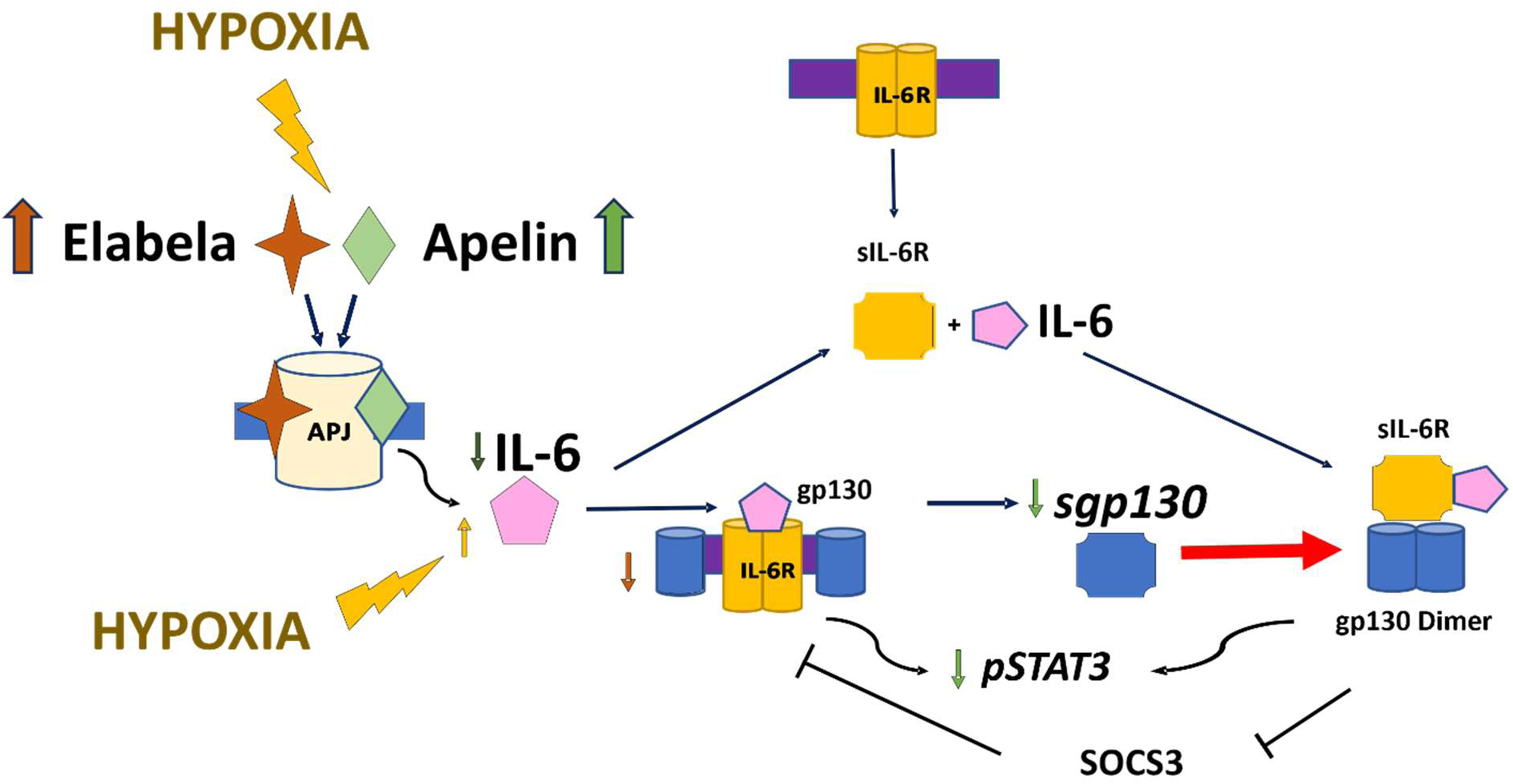

## 1. Introduction

Preeclampsia is a multisystem disorder characterized by new-onset hypertension and end-organ dysfunction with or without proteinuria [1]. Preeclampsia is a leading cause of maternal and neonatal morbidity and mortality worldwide, yet the etiology and cellular mechanisms underlying its development remain unclear. However, an imbalance of pro- and anti-inflammatory cytokines may contribute to the development of preeclampsia.

Interleukin-6 (IL-6) is a multi-functional glycoprotein that is widely expressed in gestational tissues. As both a pro- and anti-inflammatory cytokine during pregnancy, IL-6 signals through the gp130 receptor in conjunction with the IL-6 receptor (IL-6R) to modulate embryonic implantation and placental development [2, 3]. Both excess and deficiency of IL-6 have been associated with pregnancy complications including preeclampsia, fetal growth restriction, and preterm birth. For example, plasma IL-6 levels in preeclamptic women have been shown to increase after 36 weeks of gestation [4]. Endogenous IL-6 levels are partially dependent on upstream bioavailability of tumor necrosis factor-α (TNF-α), a potent inducer of further cytokine secretion [5]. Furthermore, the IL- 6/JAK/STAT3 signaling axis is modulated by levels of membrane bound and soluble IL-6 receptor availability. IL-6 receptors (IL-6R and gp130) are mediators of inflammation; thus, decreased expression of IL-6 signaling components has been shown to provide anti-inflammatory protection in preeclamptic maternal vascular systems [6]. In preeclamptic samples, maternal serum sgp130 is upregulated, while membrane-bound gp130 expression is downregulated in vascular endothelium [6]. These trends suggest that fluctuations in the components of the IL-6 receptor system might influence downstream JAK/STAT3 signaling. In particular, a recent study has shown that STAT3 expression was reduced in preeclamptic placenta, potentially contributing to the pathophysiology of the early-onset form of the preeclampsia with inefficient trophoblast migration, invasion, and proliferation [7].

The apelin receptor (APJ) is a G-protein coupled receptor with diverse downstream effects that are expressed in a variety of tissues. Two known ligands of APJ, apelin and elabela, are involved in the regulation of angiogenesis, vasodilation, inflammation, and cell survival. Both apelin and elabela are found in a variety of tissues, including the placenta. Multiple forms of apelin and elabela are derived from their respective pro-peptide precursors. Earlier work in this field has shown that systemic activation of the APJ system has both cardio- and renoprotective effects in models of heart failure and atherosclerosis [8]. Apelin is upregulated by hypoxia in vascular endothelial and other cell types potentially through hypoxia-inducible factor-1 (HIF-1) [9, 10]. Previous studies have demonstrated the expression of apelin, elabela, and APJ in human syncytiotrophoblasts, cytotrophoblasts, and endothelial cells of fetal capillaries in normal and preeclamptic placentas at term [11–14]. Our group was first to report that total apelin content, determined by quantitative radioimmunoassay, was lower in human preeclamptic chorionic villi versus normal placenta [13]. In addition, we showed that systemic administration of (Pyr^1^)-apelin-13 at late gestation reduced blood pressure and proteinuria in rats with preeclamptic features, suggesting potential cardio- and renoprotective effects of apelin in preeclampsia [15]. Furthermore, reduced expression of shorter forms of apelin in human preeclamptic versus normal chorionic villi suggested less influence of apelin on the regulation of fetal-maternal interface in preeclampsia [13]. However, elabela mRNA and protein levels were unchanged in the placenta of preeclamptic women versus normotensive controls [16]. In addition, the elabela deficiency was associated with the preeclamptic phenotype which was reversed by the administration of the elabela peptide [16, 17].

The mechanisms underlying the role of the apelin receptor axis in the placenta are not well defined. Considering the importance of the uteroplacental interface for the development of preeclampsia, the goal of this study was to determine the response of APJ system to a hypoxic-like environment and identify the actions of apelin and elabela in hypoxic HTR-8/SVneo trophoblast cells. Two ligands of the APJ receptor, (Pyr1)-apelin-13 and elabela-32, were used in the study [18]. HTR-8/SVneo cells were treated with 200 µM CoCl_2_ to mimic hypoxic conditions. Previous studies have used this method to mimic hypoxic conditions in HTR-8/SVneo cells [19, 20]. Our findings demonstrate that CoCl_2_-induced interleukin-6 (IL-6) signaling is partially downregulated by apelin and elabela in HTR-8/SVneo cells. These data expand our understanding of the protective actions of APJ signaling in preeclampsia where placental hypoxia/ischemia contributes maternal syndrome.

## 2. Materials and methods

### 2.1. Cell culture

HTR-8/SVneo cells were a kind gift from Dr. Charles Graham (Queen’s University, Ontario, Canada). The human HTR-8/SVneo cell line is a transformed first trimester human extravillous trophoblast cell line of unspecified sex that was isolated from the placenta during the first trimester of pregnancy [38]. The cell line was maintained in RPMI medium (ThermoFisher Scientific Inc., Waltham, MA, USA, cat. No. 11875093) containing 5% fetal bovine serum (FBS). To mimic hypoxia, cobalt chloride (CoCl_2_), (Sigma-Aldrich, St. Louis, MO, USA; cat. No. 7791-13-1) was added to cell culture dishes at a concentration of 200 µM. CoCl_2_-induced chemical hypoxia is a commonly used model to mimic hypoxia in cells. The exposure to CoCl_2_ results in stabilization of hypoxia-inducible factors 1α and 2α under normoxic cell culture conditions that last for several hours [39].

HTR-8/SVneo cells were exposed to apelin-13 ((Pyr-1)-Apelin-13, cat. No. AS-60833; Anaspec Inc., Fremont, CA, USA) or elabela-32 ([pGlu1]-ELA-32 (Human); cat. No. 007-25; Phoenix Pharmaceuticals Inc., Burlingame, CA, USA) at 10^-10^ M to 10^-6^ M concentrations for 24 hours. After collection of the conditioned cell culture media, the cells were washed with 1X Gibco dPBS (Fisher Scientific, Waltham, MA, USA) and collected after 20-minute incubation on ice with lysis buffer containing 10 mM Hepes/125 mM NaCl, pH 7.4, 10 µL/mL, protease inhibitor cocktail (Sigma-Aldrich, St. Louis, MO, USA; cat. No. P8340), 10 µL/mL phosphatase inhibitor cocktail 2 (Sigma-Aldrich, St. Louis, MO, USA; cat. No. P5726), 1 mM EDTA, and 1% Triton X-100. Both the cell lysate and conditioned media were frozen at -80℃ after centrifugation for future analysis of protein levels.

### 2.2. Apelin and elabela protein quantification by ELISA

HTR-8/SVneo cells were seeded at a density of 1x10^6^ cells per 100 mm dishes and grown to ∼80% confluency. CoCl_2_ was added to cells at a 200 µM concentration in 1% FBS/RPMI media, with control cells exposed to 1% FBS/RPMI media. ELISA kits were used for apelin (RayBiotech Inc., Norcross, GA, USA, cat. No. EIA-APC) and elabela (Phoenix Pharmaceuticals Inc., Burlingame, CA, USA; cat. No. EK-0017-19) protein quantification in media and cell supernatant per manufacturer’s recommended protocol.

### 2.3. Total RNA isolation and quantitative real-time RT-PCR

HTR-8/SVneo cells were seeded at a density of 1x10^5^ cells per well in 6-well culture plates and grown to ∼80% confluency. After 2-hour incubation in 1% FBS medium, CoCl_2_ was added to induce chemical hypoxia at a 200 µM concentration. Cells were also exposed to apelin or elabela at a range of concentrations from 10^-10^-10^-6^ M. Control cells were exposed to 1% FBS medium. After 1-24 hours, total RNA was extracted from the cells with the Rneasy Mini Kit (QIAGEN, Hilden, Germany) according to the manufacturer’s protocol. Rnase-free Dnase I (QIAGEN, Hilden, Germany) was used to eliminate possible DNA contamination from cells during total RNA extraction and purification. The concentration and purity of total RNA were measured with a DS-11 FX+ Spectrophotometer (DeNovix Inc, Wilmington, DE). First-strand cDNA was synthesized from 2 µg of total RNA by reverse transcription using the Omniscript RT Kit (QIAGEN, Hilden, Germany) and oligo(dT)_16_ primers (ThermoFisher Scientific Inc., Waltham, MA). The primer sequences used in the forward and reverse direction are shown in Table 1. Quantitative real-time RT-PCR was performed using the QuantStudio 3 Real-Time PCR System (Applied Biosystems, Foster City, CA). Gene specific oligonucleotide primers were used based on the published reports (Table 1). Each reaction consisted of 6 µl of 1:50 diluted first-strand cDNA, 12.5 µl of 2x RT^2^ SYBR Green qPCR Mastermix (QIAGEN, Hilden, Germany) and 0.2 µM of forward and reverse primers for a total volume of 25 µl. The PCR protocol was comprised of one cycle of 95°C for 10 min, followed by 40 cycles of 95°C for 20 sec, 60°C for 45 sec, and 72°C for 30 sec. After the reaction was completed, a melting curve analysis was performed. All samples were run in duplicates and β- actin was chosen as the reference gene for this study. The relative target mRNA levels in each sample were normalized to β-actin in the same sample using the 2^-ΔΔCt^ method (Ct = threshold cycle) and reported relative to the mean value of the control group.

**Table 1.**
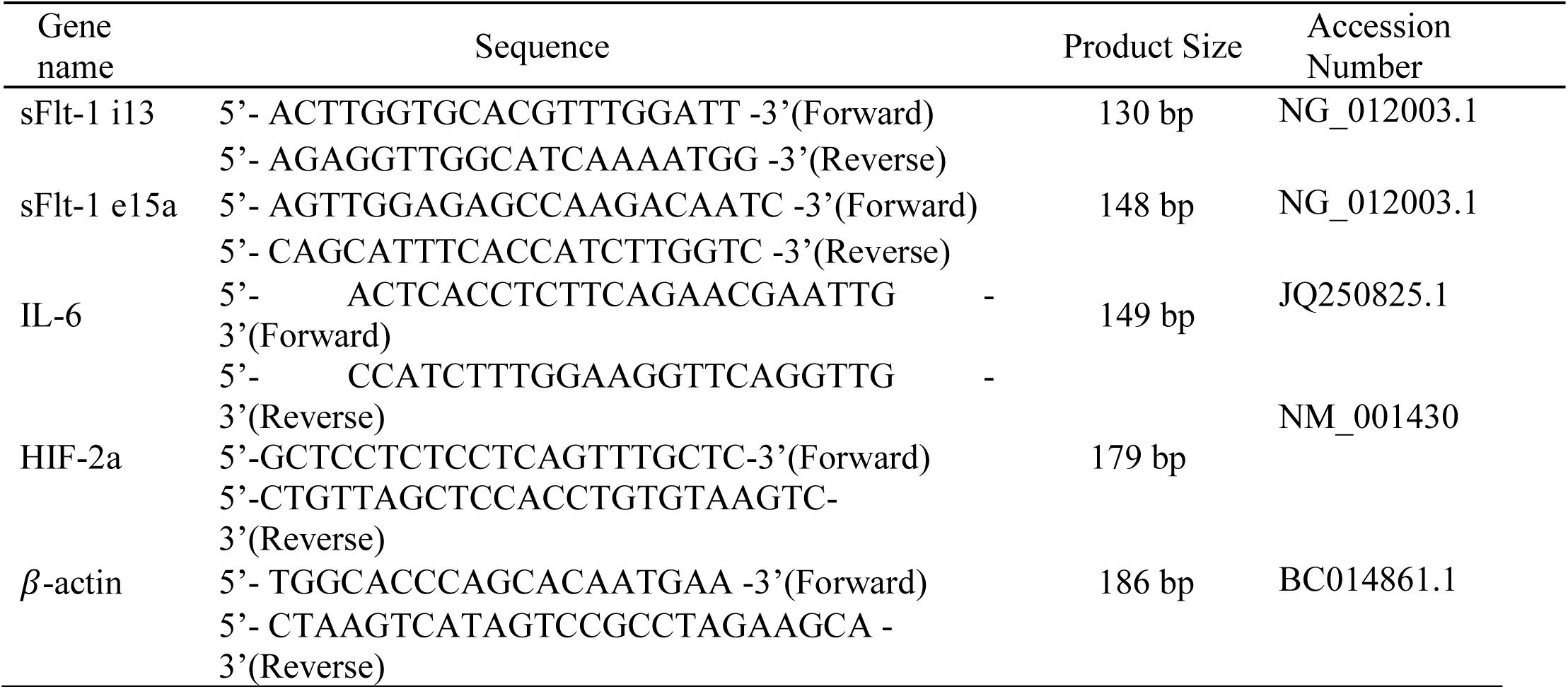
Human primer sequences for real-time RT-PCR. The primer sequences for sFlt-1 il3, sFlt-1 e15a, IL-6, HIF-2a, and *β*-actin were used for quantitative real time RT-PCR.

### 2.4. Western blotting for APJ receptor, gp130, IL-6R, SOCS-3, pSTAT3, STAT3

HTR-8/SVneo cells were collected using lysis buffer as described above. The collected cell lysate samples were subjected to 3-second sonication pulses at 20 kHz for 15 seconds. Samples were then centrifuged (10,000 g, 5 min, 4℃) to remove debris. Total protein levels in the supernatant were determined using the Pierce BCA Protein assay kit (ThermoFisher Scientific Inc., Waltham, MA, USA) according to the manufacturer’s protocol. 15µg of protein from each sample were separated by 4-20% TGX Stain-Free gel (Bio-Rad Laboratories, Cat#: 567-8095) and transferred onto a PVDF membrane with the Trans-Blot Turbo Transfer System (Bio-Rad Laboratories). The membrane was blocked with 5% non-fat milk or BSA in Tris-buffered saline with 0.05% Tween (blocking buffer) for 1 hour at room temperature, then incubated with primary antibodies in the blocking buffer overnight at 4°C. Following incubation with the corresponding horseradish peroxidase-conjugated secondary antibody at room temperature for 1 hour, protein bands were visualized by using SuperSignal^TM^ West Femto Kit (Thermo Scientific, Cat #: 34096). The images were captured using a ChemiDoc Imaging System (Bio-Rad Laboratories) and analyzed using Image Lab software (Bio-Rad Laboratories). Each band was expressed in arbitrary units and normalized to β-actin. The primary antibodies used were rabbit anti-β-actin antibody (1:1000 dilution; Cell Signaling Technology, Danvers, MA; Cat #: 8457), rabbit anti-APJ antibody (1:500 dilution; MilliporeSigma, Burlington, MA; Cat #: ABD43), rabbit anti-gp130 antibody (1:500 dilution; Cell Signaling Technology, Danvers, MA; Cat #: 3732), mouse anti-IL6Rα antibody (1:100 dilution; *Santa Cruz Biotechnology*, Dallas, TX; Cat #: sc-373708), mouse anti-SOCS3 antibody (1:100 dilution; *Santa Cruz Biotechnology*, Dallas, TX; Cat #: sc-73045), mouse anti-STAT3 antibody (1:500 dilution; Cell Signaling Technology, Danvers, MA; Cat #: 9139), rabbit anti-pSTAT3 antibody (1:500 dilution; Cell Signaling Technology, Danvers, MA; Cat #: 9145).

### 2.5. IL-6, soluble IL-6R (sIL-6R), and soluble gp130 (sgp-130) Enzyme-Linked Immunosorbent Assays (ELISA) in conditioned media

Total protein levels were determined in the conditioned media following apelin and elabela treatments in HTR-8/SVneo cells using the Pierce BCA Protein assay kit (ThermoFisher Scientific Inc., Waltham, MA, USA) according to the manufacturer’s protocol. Individual ELISA kits were used for IL-6 (RayBiotech Inc., Norcross, GA, USA; cat. No. ELH-IL6), sIL-6R (RayBiotech Inc., Norcross, GA, USA; cat. No. ELH-IL6sR), and sgp130 (RayBiotech Inc., Norcross, GA, USA; cat. No. ELH-sgp130) protein quantification in the conditioned media 24 hours after treatment.

### 2.6. Statistical Analyses

Study groups were compared using one-way analysis of variance (ANOVA) with- (Figure 1) or without repeated (Figures 3-7) measures followed by the Bonferroni post-hoc tests (GraphPad 10 Software, San Diego, CA). For Figure 2, the comparisons between studied groups were made using unpaired t-test. A p-value of less than 0.05 was considered statistically significantly different. All data were presented as mean ± SEM.

**Figure 1.**
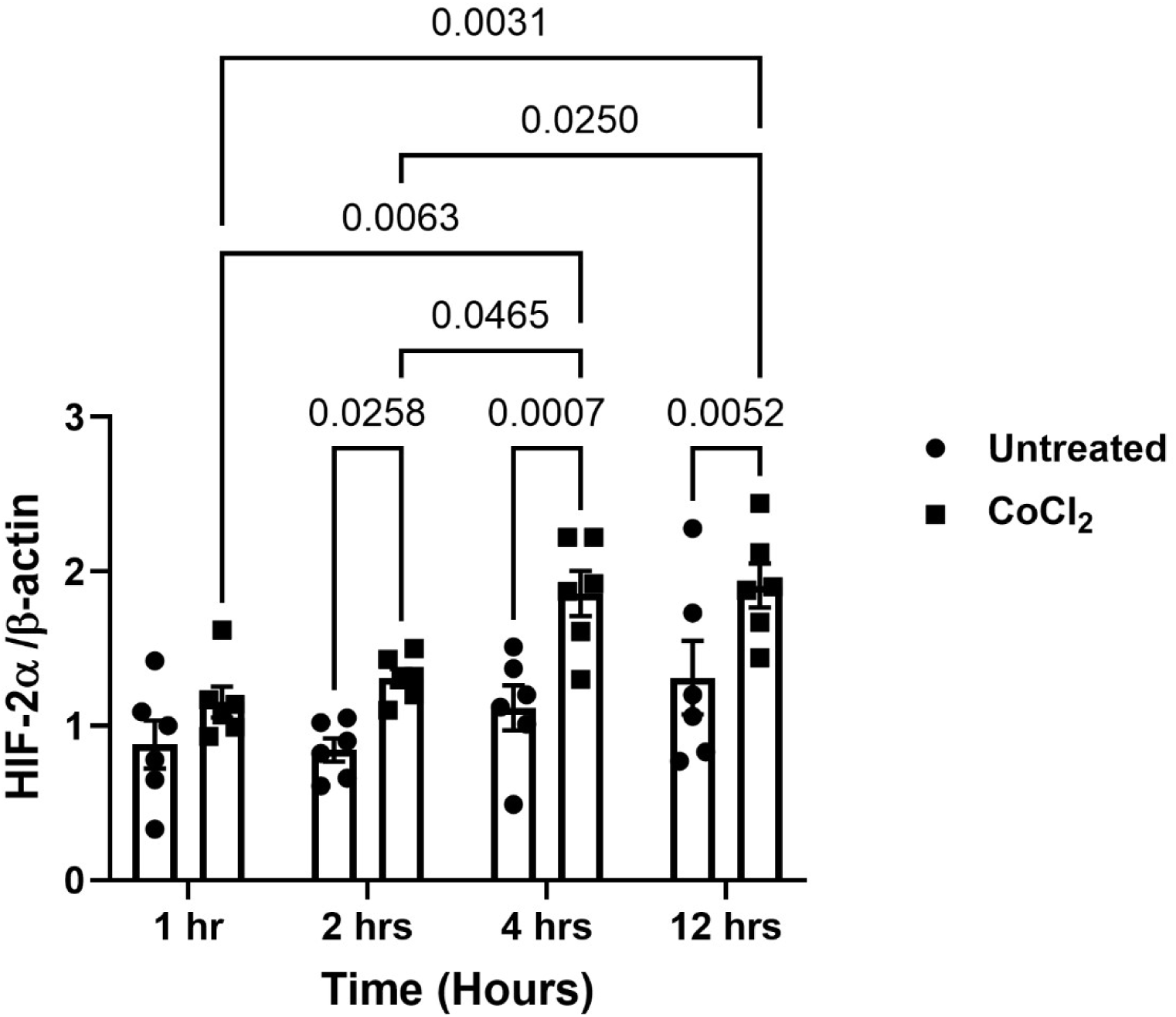
A time-dependent upregulation of HIF-2α in response to CoCl_2_ in HTR-8/SVneo cells. Data are mean ± SEM, n=6.

**Figure 2.**
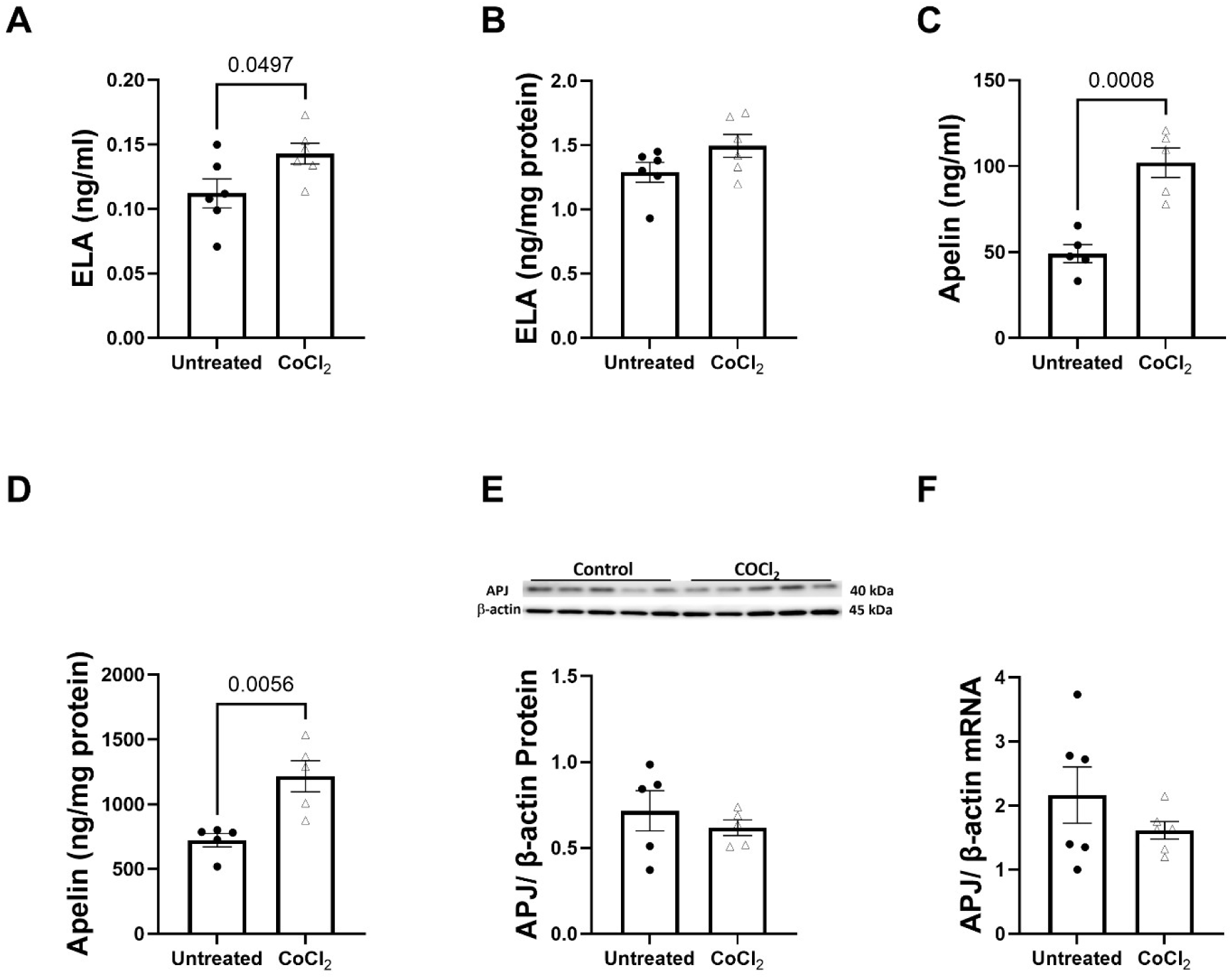
The effect of CoCl_2_ on the levels of elabela (ELA), apelin, and apelin receptor APJ. The data were obtained using ELA or apelin ELISA assays in HTR-8/SVneo cell homogenates (Panels A and C) or conditioned media (Panels B and D), western blotting (Panel E), or real-time RT-PCR (Panel F) for APJ. Data are mean ± SEM, n=5-6.

**Figure 3.**
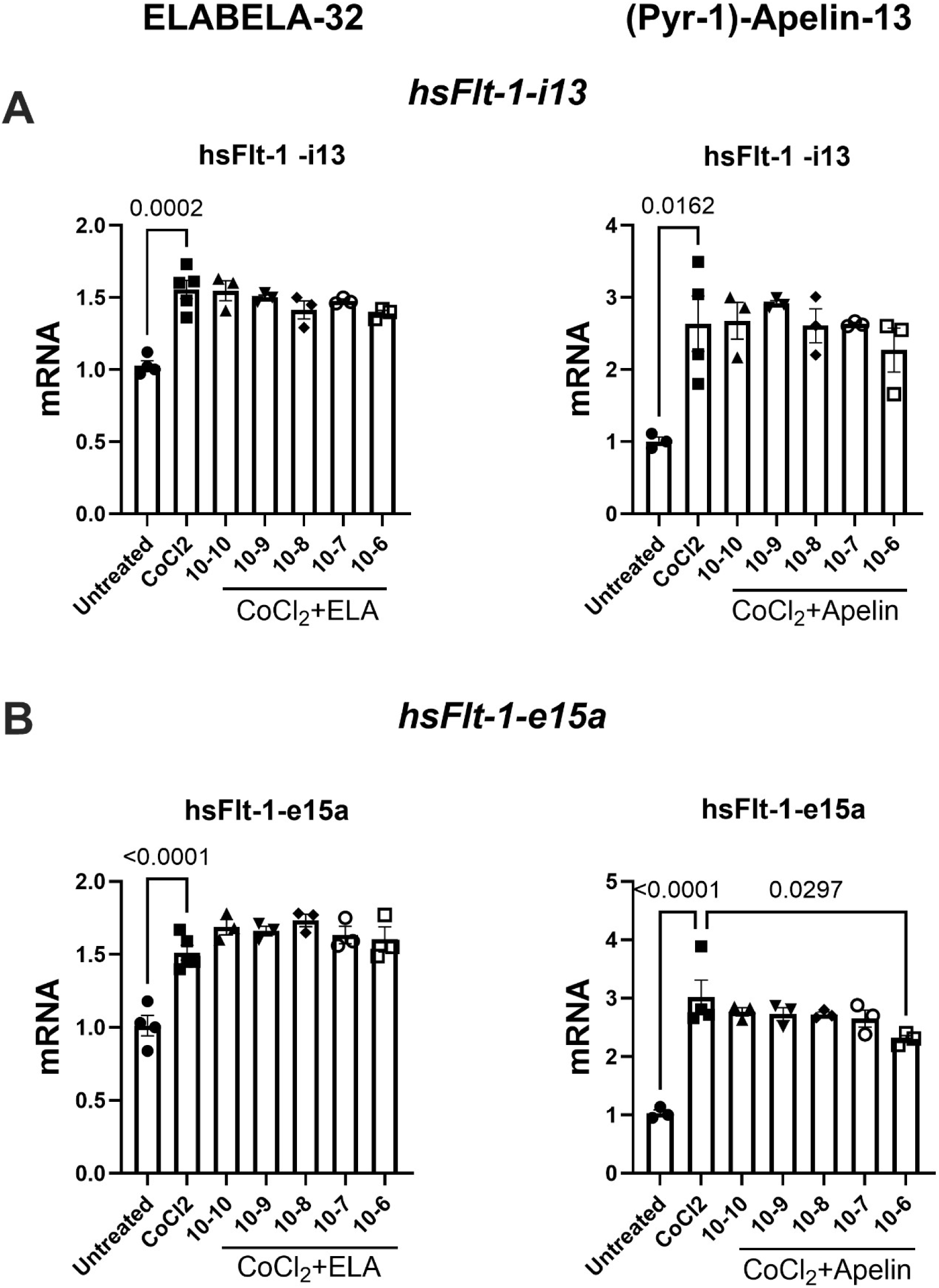
Elabela (ELA) or apelin treatment did not alter sFlt-1 variants in CoCl_2_-treated HTR- 8/SVneo cells. Data are mean ± SEM, n=3-6.

## 3. Results

### 3.1. The exposure to CoCl_2_ increased elabela and apelin protein levels in HTR-8/SVneo cells and conditioned media (Figures 1-2)

We first establish whether the exposure to CoCl_2_ induces the expression of one of the hypoxia-responsive genes HIF-2α in HTR-8/SVneo cells. Figure 1 demonstrates a time-dependent upregulation of HIF-2α in response to CoCl_2_ in HTR-8/SVneo cells. To examine the endogenous response of the APJ axis to hypoxia, we quantified apelin and elabela content by ELISA after 24-hour incubation with CoCl_2_. Figure 2A and 2B show that CoCl_2_ increased total levels of elabela in the conditioned media, but not in the cell supernatant of HTR-8/SVneo cells. However, apelin content was increased approximately 2-fold in both the cell lysate and conditioned media of CoCl_2_-exposed cells (Figure 2C-D). Elabela levels also were increased but were 100-fold less when compared to the increase in apelin content. We also quantified APJ receptor mRNA and protein levels in CoCl_2_-treated HTR-8/SVneo cells, but CoCl_2_ treatment did not alter the levels of APJ receptor (Figure 2E-F).

### 3.2. The effects of elabela and apelin on sFlt-1 isoforms in CoCl_2_-exposed HTR-8/SVneo cells (Figure 3)

To confirm that CoCl_2_ -induced hypoxia upregulated sFlt-1 and establish the effect of apelin and elabela on sFlt-1 expression, the cells were treated with a range of apelin or elabela peptide concentrations while being exposed to CoCl_2._ The exposure to CoCl_2_ increased the mRNA levels of sFlt-1-1-i13 (2.6-fold) and sFlt-1-1-e15a (2.9-fold); however, neither apelin nor elabela treatment affected sFlt-1-variants at the physiologically relevant concentrations (<10^-7^M).

### 3.3. Both elabela and apelin attenuated the levels of IL-6 mRNA in CoCl_2_-exposed HTR- 8/SVneo cells (Figure 4)

CoCl_2_ upregulated IL-6 mRNA levels (Figure 4A-B) while treatment with elabela (0.1 μM) or apelin downregulated the cellular levels of IL-6 mRNA (Figures 4A and B, respectively).

**Figure 4.**
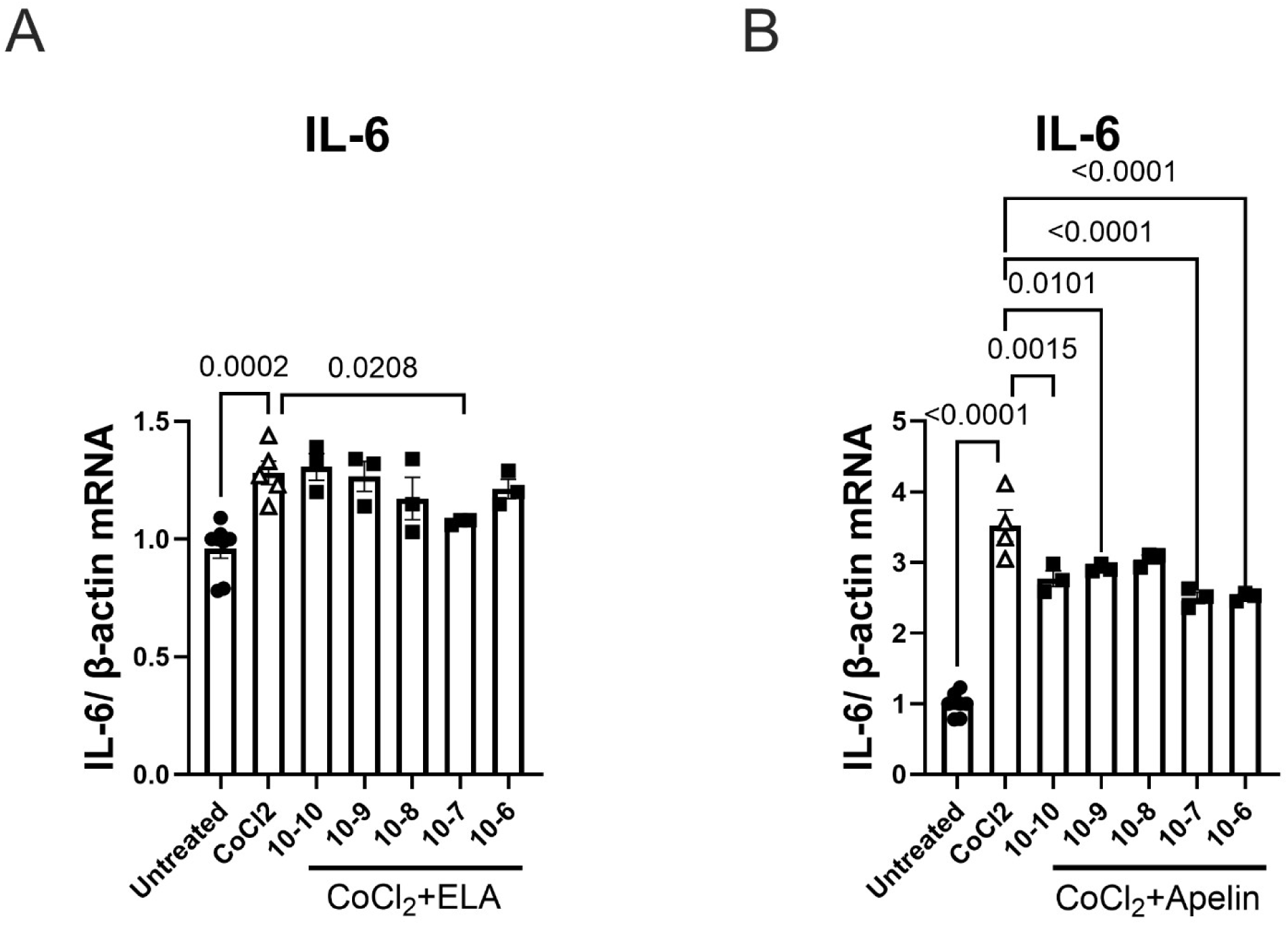
Elabela (ELA) or apelin treatment decreased IL-6 mRNA in CoCl_2_-treated HTR-8/SVneo cells. Data are mean ± SEM, n=3-6.

### 3.4. Apelin or elabela effect on soluble and membrane forms of gp130 in CoCl_2_-exposed HTR- 8/SVneo cells (Figure 5)

Soluble gp130 (sgp130) protein levels were upregulated by CoCl_2_ (Figure 5A, C). A low concentration of apelin decreased sgp130 levels in CoCl_2_-exposed cells (Figure 5A). In contrast, elabela had no effect on sgp130 protein levels in HTR-8/SVneo cells. (Figure 5C). Figure 5B shows that apelin did not alter the levels of membrane-bound gp130 protein, however elabela increased gp130 at both studied concentrations (Figure 5D).

**Figure 5.**
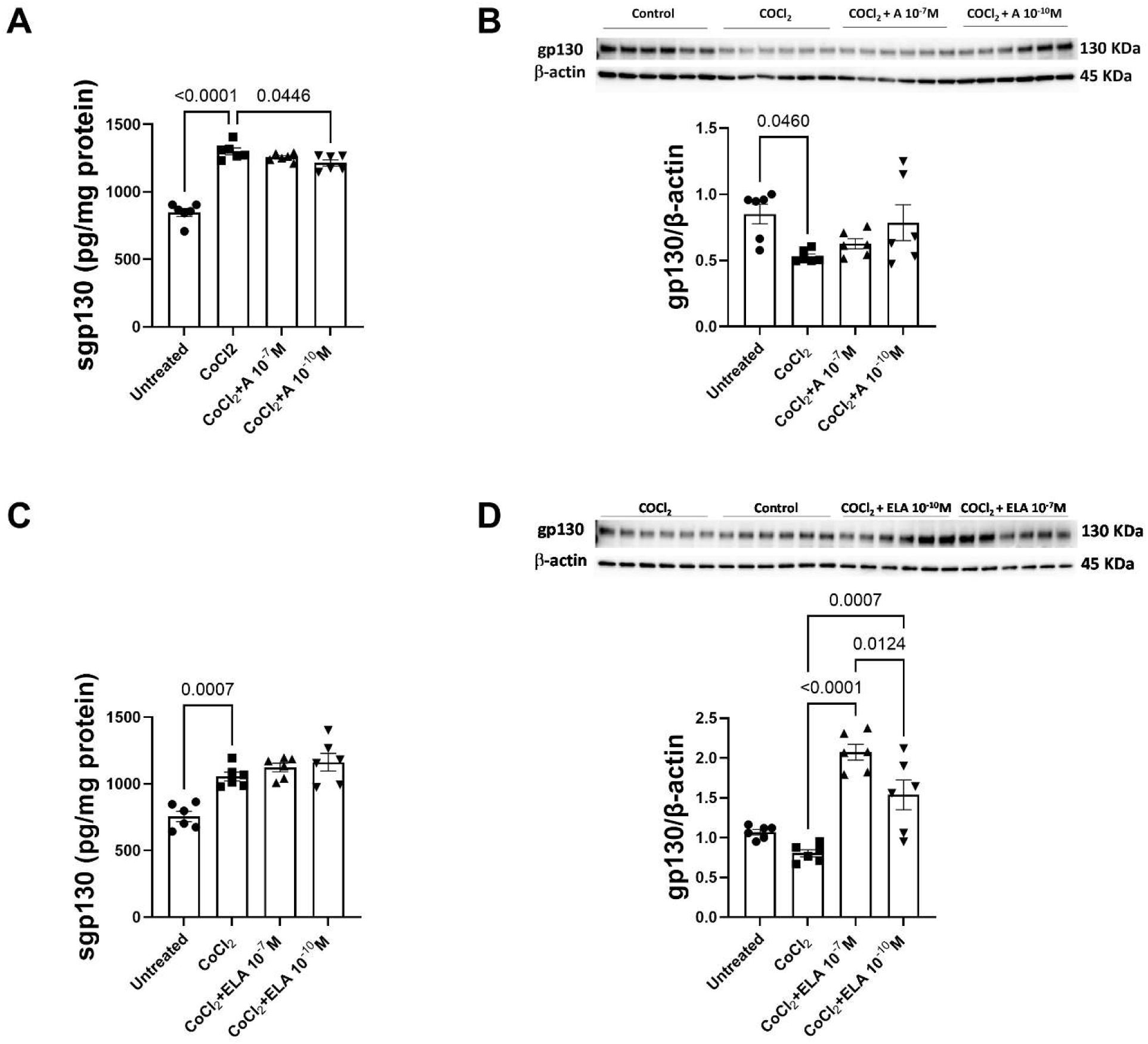
Soluble and membrane-bound gp130 in response to apelin (A; Panels A-B) or elabela (ELA; Panels C-D) in CoCl_2_-treated HTR-8/SVneo cells. Data are mean ± SEM, n=6 in each group.

### 3.5. Apelin and elabela effects on membrane- and soluble IL-6R in CoCl_2_-exposed HTR- 8/SVneo cells (Figure 6)

IL-s6R concentrations were increased in CoCl_2_-exposed cells (Figure 6A, C). Neither apelin nor elabela affected IL-s6R levels in CoCl_2_-exposed cells. However, elabela decreased membrane form of IL-6R levels in CoCl_2_-exposed cells at both 10^-7^ M and 10^-10^ M concentrations (Figure 6D).

**Figure 6.**
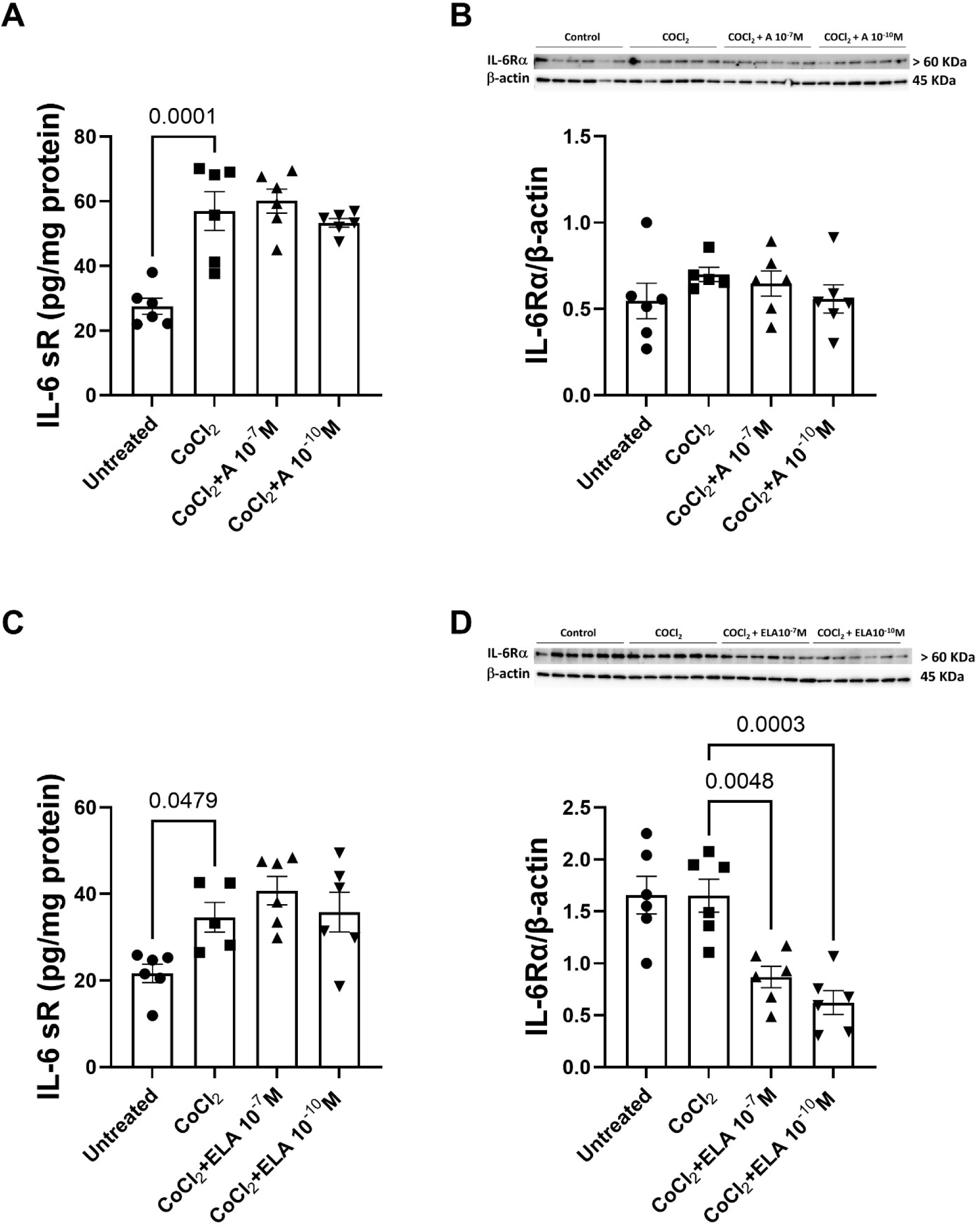
Soluble and membrane-bound IL-6R in response to apelin (A; Panels A-B) or elabela (ELA; Panels C-D) in CoCl_2_-treated HTR-8/SVneo cells. Data are mean ± SEM, n=5-6.

**Figure 7.**
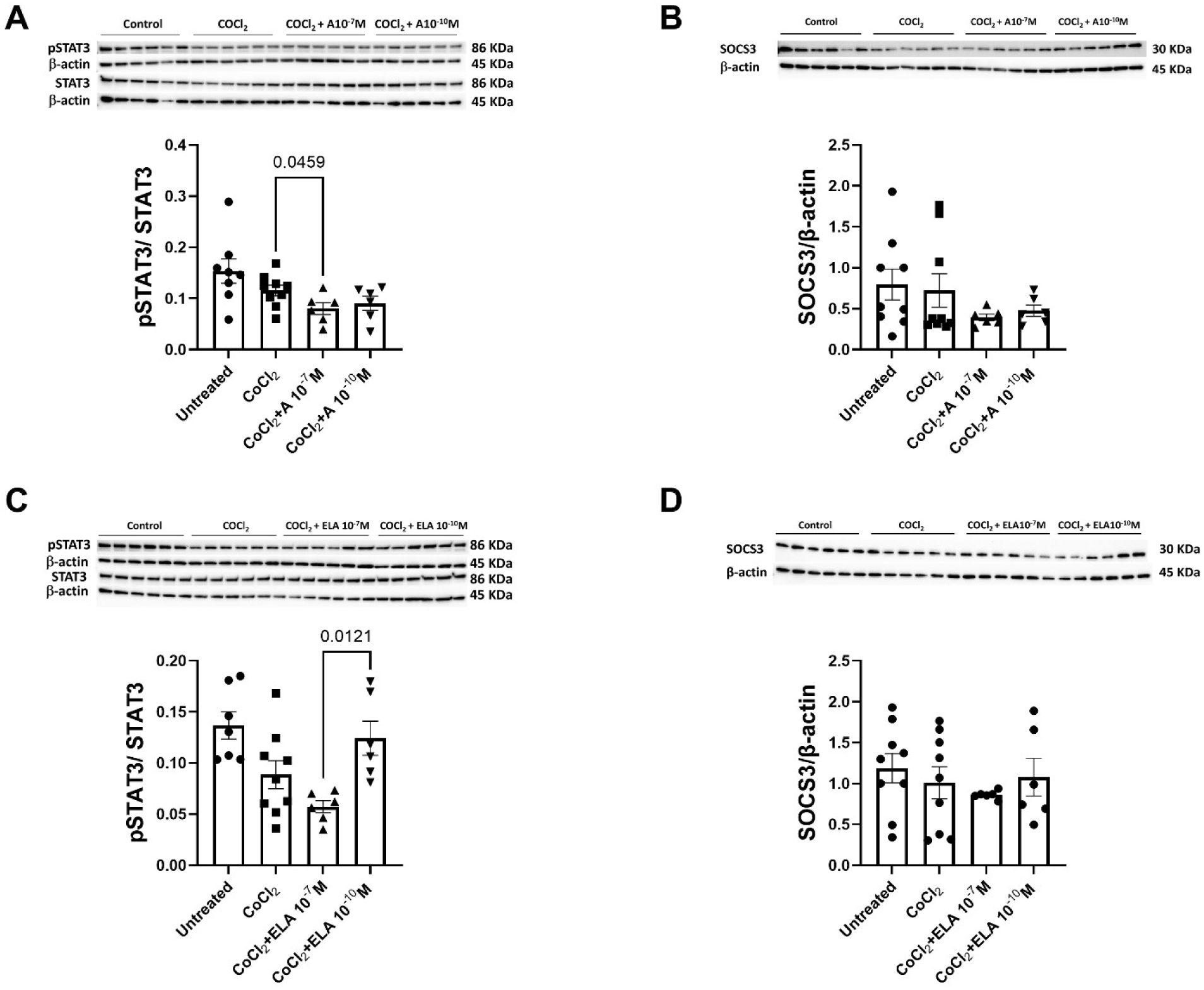
pSTAT3/STAT3 ratio and SOCS3 protein levels in response to apelin (A; Panels A and B) or elabela (ELA; Panels C and D) in CoCl_2_-treated HTR-8/SVneo Cells. Data are mean ± SEM, n=5-9.

### 3.6. pSTAT3/STAT3 and SOCS3 in CoCl_2_-exposed HTR-8/SVneo Cells (Figure 7)

No differences in the pSTAT3/STAT3 ratio were found in CoCl_2_-exposed cells. Apelin decreased the pSTAT3/STAT3 ratio at 10^-7^ M (Figure 7A). SOCS3 was not changed by CoCl_2_ (Figure 7C-D). Neither apelin nor elabela affected SOCS3 levels in CoCl_2_-treated cells (Figure 7C-D).

## 4. Discussion

Preeclampsia is a pregnancy disorder that affects 2-8% of pregnancies worldwide and characterized by hypertension, proteinuria, with- or without target organ dysfunction. Preeclampsia can lead to both maternal and fetal complications, including ischemic stroke placental abruption, fetal growth restriction, and end organ damage [21]. Despite these serious complications, the pathogenesis of preeclampsia remains poorly understood. As a result, there are no effectively outlined guidelines for the prediction, prevention, or treatment of this hypertensive pregnancy disorder [21].

Previous studies have employed hypoxic trophoblast cell lines to investigate the roles of potential biomarkers and their involvement in key placental processes implicated in preeclampsia. The apelin receptor (APJ) system has emerged as a candidate for both diagnostic and therapeutic applications in preeclampsia. Prior research has shown that apelin, elabela, and APJ are expressed in human cytotrophoblasts, syncytiotrophoblasts, and fetal endothelial cells, suggesting an autocrine/paracrine role at the feto-maternal interface. Notably, apelin levels were significantly reduced in preeclamptic compared to normal term chorionic villi [13], indicating a possible association between apelin downregulation and preeclampsia. In contrast, APJ receptor levels were comparable between the preeclamptic and normal pregnant chorionic villi at term suggesting that altered apelin concentration or receptor affinity rather than changes in the levels of the receptor account for an influence of the system in preeclampsia [13]. Our previous preclinical studies also demonstrated anti-hypertensive and renoprotective effects of apelin in a preeclamptic rat model at late gestation, suggesting a therapeutic potential of the APJ axis in preeclampsia [15]. Similarly, elabela administration in elabela knockout mice reduced blood pressure, proteinuria, and improved fetal weight at late gestation [16]. However, the role of the APJ axis in early pregnancy remains poorly understood, particularly in hypoxic environment associated with early stages of pregnancy.

It is likely that the regulation of apelin, elabela, and APJ is multifaceted under hypoxic conditions. In a subset of preeclamptic placentas, abnormal trophoblast migration creates hypoxic environment for placental cells at late gestation [21]. Studies using human HTR-8/SVneo trophoblast cells under hypoxic conditions have shown that apelin reduces apoptosis by downregulating active caspase-3 and increasing the Bcl-2/Bax ratio, while also decreasing superoxide radical production [16]. Potential pathways for apelin’s function in placental trophoblasts include PI3K/Akt, ERK1/2, STAT3, and AMPK kinase, which can be regulated via the APJ receptor system [22, 23]. Hypoxic conditions have been shown to reduce the expression of signaling kinases PI3K and Akt levels [22], thus potentially leading to the endothelial cell dysfunction seen in preeclampsia. However, hypoxia has also been shown to increase expression of the APJ receptor and as reported in our study, promotes apelin and elabela upregulation [22, 23]. Additionally, apelin and elabela have been reported to increase trophoblast proliferation, migration, and invasion through the activation of MAPK/ERK signaling [24–26]. These peptides may counteract hypoxia-induced reductions in PI3K and Akt expression, thereby improving trophoblast function [16]. Thus, the goal of this study was to examine the response of HTR-8/SVneo trophoblastic cells to exogenous apelin or elabela focusing on pro-inflammatory cytokine signaling in the setting of CoCl_2_-induced model of hypoxia. We found that CoCl_2_- upregulated apelin and elabela levels. Local activation of this peptidergic system may be a compensatory response of the HTR-8/SVneo cells to hypoxia as exogenous apelin and elabela treatment downregulated the components of CoCl_2_-induced IL-6 signaling.

We found that CoCl_2_ increased endogenous apelin and elabela peptide levels in HTR-8/SVneo cells without changing the concentration of the APJ receptor. This response is consistent with other studies that have shown enhanced apelin expression under hypoxic conditions in cultured cardiomyocytes and endothelial cells [27]. In hypoxia-exposed cardiac endothelial cells, apelin levels were increased when compared to cells exposed to control oxygen levels [27]. Apelin promoted further angiogenesis and mobilization of endothelial cell progenitors following ischemia induced conditions [28]. Previous work using BeWo and JEG-3 trophoblast cell lines has demonstrated a potential role for apelin in regulating the G2/M phase of the cell cycle thus promoting cell survival of trophoblast cells [23]. While the exact mechanisms are unknown, one potential mechanism of apelin or elabela upregulation in response to CoCl_2_-induced model of hypoxia is the binding of HIF-1α to a hypoxia-responsive element (HRE) on the first intron of the peptide gene. Although, not tested in this study, the upregulation of apelin or elabela in response to CoCl_2_-induced hypoxia may potentially be a compensatory response to increase the survival of trophoblast cells.

The exposure to CoCl_2_ markedly increased levels of IL-6 mRNA in HTR-8/SVneo cells. Appropriate regulation of pro- and anti-inflammatory cytokines is necessary for healthy placentation and pregnancy progression. However, altered cytokine expression can lead to abnormal placentation and subsequent complications including preeclampsia. As a multifunctional cytokine, IL-6 is important for the regulation of early implantation events, however an excessive IL-6 bioavailability may potentially be harmful for the placentation by inhibiting the generation of CD4+ T regulatory cells required for pregnancy tolerance [3]. The addition of apelin or elabela at physiologically relevant concentrations significantly decreased IL-6 mRNA levels. This finding agrees with previous studies demonstrating that apelin downregulated IL-6 and TNF-α mRNA levels in pulmonary vascular endothelial cells [23]. A possible mechanism for apelin to reduce IL-6 concentrations is through the regulation of the upstream signaling such as NF-kB or counteracting the actions of miR- 15b-5p [29]. Additional studies are in progress to investigate the mechanisms of apelin or elabela actions on IL-6 and other proinflammatory cytokines in trophoblast cells in response to direct exposure to low oxygen environment.

Additionally, we show that mRNA levels of sFlt-variants were significantly increased under hypoxic conditions. sFlt-1 and sEng are proteins with antiangiogenic properties. In an environment of ischemia and hypoperfusion, as seen in preeclampsia, sFlt-1 is upregulated leading to further vasoconstriction and vascular endothelial dysfunction [22, 30]. sFlt-1 also serves as an important biomarker for vascular dysfunction in preeclamptic placentas. While hypoxic conditions produced a significant increase in sFlt-1 mRNA in our study, neither apelin nor elabela had an effect on sFlt-1 variants apart from apelin downregulating sFlt-1 mRNA at a supraphysiological concentration. These data agree with our previous study where systemic administration of (Pyr^1^)-apelin-13 did not alter circulating sFlt-1 levels at late gestation in preeclamptic rats [15]. These findings suggest that apelin and elabela modulate the hypoxic response of HTR-8/SVneo cells through mechanisms independent of sFlt-1 signaling. However, previous research has shown that apelin can significantly decrease serum sFlt-1, sEng, and IFN-y while increasing the serum levels of VEGF and PLGF in a RUPP model of preeclampsia [30]. It is possible that the effect of apelin on sFlt-1 is model-specific. In addition, apelin can influence other components of pro- and anti-angiogenic pathways such as promoting angiogenesis through the upregulation of VEGF and PLGF via APJ/PI3K/ERK pathway [30].

A pro-inflammatory IL-6 trans-signaling is known to be highly regulated by sgp130 as opposed to the membrane-bound form of the receptor [31]. Furthermore, sgp130 levels are increased in the plasma of preeclamptic women [6]. Thus, we focused on the effect of apelin or elabela on the soluble gp130 protein levels in CoCl_2_-treated cells. Our data show that soluble gp130 protein levels were decreased by apelin in CoCl_2_-exposed trophoblast cells. Considering that similar to apelin and elabela, IL-6 and gp130 are highly localized to syncytiotrophoblast layer of villous tissue of the placenta [32], a decrease in soluble gp130 may be a part of the overall anti-inflammatory actions of apelin on IL-6 signaling. Thus, apelin-induced downregulation of sgp130 may contribute to the reduction of pro-inflammatory milieu in hypoxic placenta.

Another potential mechanism of the relationship of apelin with IL-6/sGP130 signaling may involve IL-6/IL-6R-induced-mediated JAK mediated phosphorylation of gp130, ultimately leading to increased cytokine production [33]. Furthermore, sgp130 inhibition may upregulate the suppressor of cytokine signaling 3 protein (SOCS-3) activity [34, 35]. SOCS-3, a known inhibitor of classical IL-6 signaling, is located downstream of IL-6, and is shown to be reduced in human preeclamptic placenta [6]. However, neither apelin nor elabela had an effect of SOCS-3 in CoCl_2_- treated HTR-8/SVneo cells. It has been suggested that SOCS-3 upregulation may serve an anti-inflammatory role in hypoxic conditions by increasing production of IL-10 [34, 35]. SOCS-3 can mediate both the IL-6/gp130/JAK and IL-6/IL-6R signaling pathways, and the presence of increased levels of IL-6 produced a subsequent increase in anti-inflammatory IL-10 production in JEG-3 trophoblast cells [35]. Of note, both SOCS-3 and IL-10 levels are significantly downregulated in placental trophoblasts from preeclamptic women [35], suggesting decreased influence of these cytokines on the regulation of immunological tolerance at feto-maternal interface. Finally, the levels of soluble IL-6R were increased by CoCl_2_-induced hypoxia, however neither apelin nor elabela influenced its expression in HTR-8/SVneo cells. However, elabela downregulated the levels of membrane-bound IL-6R. While little is known about the direct effects of apelin or elabela on soluble or membrane-bound IL-6R levels, lower levels of IL-6R were detected in early onset severe PE placentas [36]. In contrast, lower levels of IL-6R have been shown to decrease proliferation and increase apoptosis suggesting a complex regulation of IL-6 signaling in trophoblast cells [36].

Limitations of the study: This study has several limitations inherent to the use of HTR-8/SVneo cells as an *in vitro* model mimicking the trophoblast response to exogenous stimuli. As an immortalized cell line, HTR-8/SVneo cells do not fully recapitulate the phenotype or gene expression patterns of primary trophoblasts *in vivo* [37]. In addition, a monoculture system used in this study lacks the cellular diversity of the human placenta. Furthermore, although the relevance of using CoCl_2_ as a model of chemical hypoxia was supported by the time-dependent upregulation of hypoxia-responsive genes such as HIF-2α and sFlt-1 in HTR-8/SVneo cells in our study, future studies will focus on the direct exposure of HTR-8/SVneo cells to low oxygen conditions to more closely mimic the effects of physiological hypoxia and increase the translatability of our findings to *in vivo* placental environment.

## 5. Conclusion

Our data confirmed that both membrane and soluble forms of IL-6R, and the components of IL-6 trans-signaling such as soluble and membrane-bound gp130 are present in HTR-8/SVneo cells. We showed that these cytokines can be modulated by CoCl_2_, and/or by apelin or elabela. Since IL-6 signaling through its soluble receptor primarily drives inflammation, whereas signaling via the membrane-bound receptor tends to support cell survival and anti-inflammatory effects, our data suggest that apelin or elabela could be involved in the complex regulation of the hypoxic environment in trophoblast cells relevant for preeclampsia.

## Abbreviations

CoCl_2,_: cobalt chloride
APJ,: apelin receptor
VEGF,: vascular endothelial growth factor
PlGF,: placental growth factor
TNF-α,: tumor necrosis factor alpha
IL-6,: interleukin-6
IL-6R,: interleukin-6 receptor
sIL-6R,: soluble interleukin-6 receptor
sFlt,: soluble fms-like tyrosine kinase
gp130,: glycoprotein 130
associated with IL-6R sgp130,: soluble glycoprotein 130
JAK2,: Janus kinase 2
non-receptor tyrosine kinase STAT3,: signal transducer and activator of transcription
3 HIF-1 α,: hypoxia-inducible factor 1α
SOCS-3,: suppressor of cytokine secretion 3

## Data availability

All data supporting this study is presented in the Methods and Results section of this manuscript.

## Credit authorship contribution statement

Anna Soloshenko - Data curation; Formal analysis, Writing – review & editing, Writing – original draft, revisions, Conceptualization

Courtney Brown – Writing – review & editing, Writing – original draft, revisions

Xuming Sun - Data curation; Formal analysis, Writing – review & editing, Writing – original draft, revisions, Conceptualization

Ajay N. Roy – Data curation, Formal analysis

Jonathan Ray – Writing – review & editing, Writing – original draft, revisions Mark C. Chappell - Writing – review & editing, Writing – original draft, revisions

Liliya M. Yamaleyeva - Writing – review & editing, Writing – original draft, Supervision, Project administration, Funding acquisition, Conceptualization

## Declaration of Generative AI and AI-assisted technologies in the writing process

The authors acknowledge that no Generative AI and AI-assisted technologies were used during the preparation of this work.

